# Alcohol Increases Aggression in Flies

**DOI:** 10.1101/685529

**Authors:** Annie Park, Tracy Tran, Elizabeth Scheuermann, Linda Gutierrez, Christopher Stojanik, Julian Plyler, Grace Thompson, Dean Smith, Nigel S. Atkinson

**Affiliations:** Department of Neuroscience and Waggoner Center for Alcohol and Addiction Research, The University of Texas at Austin, Austin, TX 78712.; Department of Pharmacology and Neuroscience, University of Texas Southwestern Medical Center, Dallas, Texas 73590-9111.

## Abstract

Alcohol-induced aggression is a destructive and widespread phenomenon, but we understand very little about the mechanisms that produce this behavior. We found that two different alcohol exposures potentiate aggression in male flies. (1) A pharmacologically relevant dose of alcohol increases aggression and decreases a goal-directed behavior in male flies. (2) In addition, the odor of alcohol itself enhances intermale aggression by potentiating olfactory signaling by cis-vaccenyl acetate (cVa), a volatile male pheromone. Characterizing these behaviors in the genetically tractable fruit fly can lead to a better understanding of the molecular correlates that regulate alcohol-induced aggression in humans and provide insights into an ethologically relevant behavior.

**One Sentence Summary:** We identified two pathways through which alcohol stimulates intermale aggression in flies; one acts by potentiating a male olfactory pheromone while the other is mediated by the systemic effects of alcohol.

## Main Text

A staggering 40% of violent offenses are committed by people under the influence of alcohol, and at least 50% of sexual assault cases are associated with alcohol consumption (*1*). Despite alcohol-induced aggression being a costly and pervasive issue, little is understood about the neurobiological mechanisms that provoke individuals to become more aggressive after consuming alcohol. To help bridge this gap in knowledge, we introduce *Drosophila melanogaster* as a model organism in which male flies become more aggressive with exposure to alcohol. There exists a substantial body of work on the effects of alcohol on fly behavior as well as a considerable amount of work on aggression in flies (*2*, *3*). However, nothing is known about the intersection between these topics, that is, whether or not alcohol modulates aggression in flies. We find that a pharmacologically relevant blood alcohol concentration (BAC) promotes male aggression. Furthermore, this alcohol dosage impairs male courtship—a goal-directed behavior (*4*). In addition, there exists a secondary mechanism through which alcohol induces aggression. We find that the mere odor of alcohol increases aggression. This olfactory response occurs in the absence of intoxication and in the absence of a detectable BAC. One of the most well-studied fly pheromones, cis-vaccenyl acetate (cVa), is a male-produced pheromone that is known to cause aggression in male flies (*5*). We find that smelling alcohol potentiates the cVa pathway and that this leads to increased aggression. This work demonstrates that *Drosophila melanogaster* is an ideal genetic model system for studying the molecular origins of alcohol-induced aggression.

## Results

### A pharmacologically relevant dose of alcohol potentiates aggression

We tested different ethanol exposure paradigms to assess their effects on intermale aggression. One hour following a 5-minute exposure to the vapor from a 30% ethanol solution (abbreviated PEA for Post-Ethanol Aggression) we observed a substantial rise in intermale aggression (Fig. 1A and fig. S1). The blood alcohol concentrations (BAC) immediately after the 5-minute alcohol vapor treatment (Low Dose) and one hour after this treatment (PEA) were 0.047 and 0.0145 mg/mL, respectively (Fig. 1C). This latter concentration is pharmacologically relevant and is roughly equivalent to the BAC produced by half a standard drink in humans.

**Figure 1.**
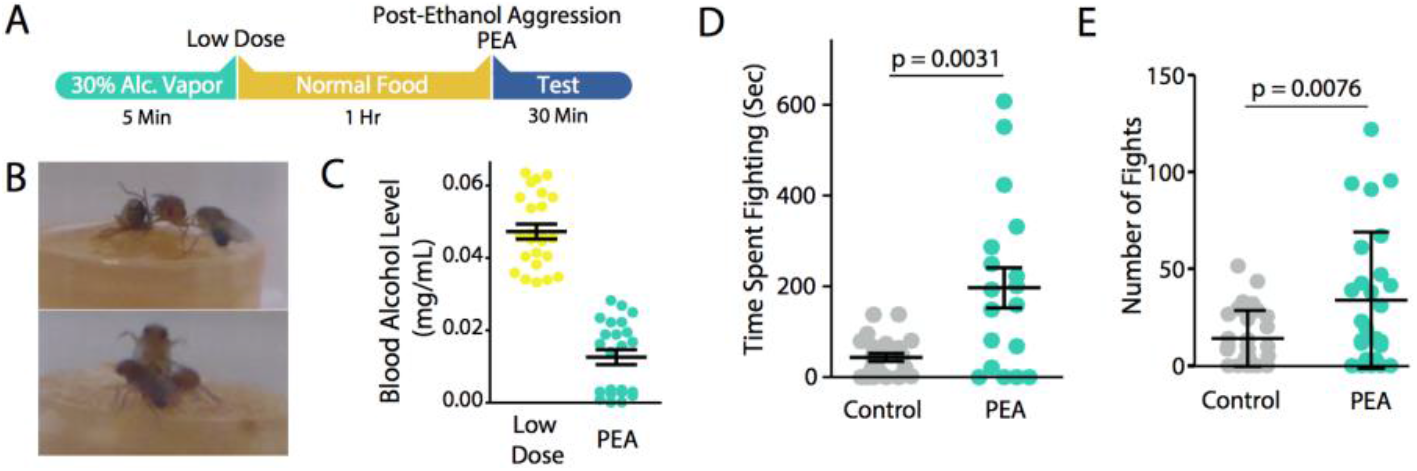
Treatment with PEA makes flies more aggressive. (**A**) Alcohol treatment protocol for Post-Ethanol Aggression. (**B**) Pictures of the flies in the aggression chamber fencing (top) and lunging (bottom). (**C**) Blood alcohol levels measured in flies immediately after alcohol exposure (Low Dose Mean ± SEM; 0.047±0.002 mg/mL, n=24) and one hour after alcohol exposure (PEA; Mean ± SEM; 0.0145±0.009 mg/mL). (**D**) Time spent fighting and (**E**) number of fights (two-tailed unpaired *t*-test, n=24 for Control and n=18 for PEA).

Video recordings were scored manually for intermale aggressive behaviors and for mating attempts with the decapitated female. Each 30-minute video recording was scored by at least two observers and averaged. In comparison to control animals, male PEA flies showed a significant increase in the time spent fighting, the number of fights they engaged in, and the number of lunges (Fig. 1, D and E; fig. S2; Movie S1 and S2). Finally, PEA males were more likely than control flies to attempt copulation with the female carcass (Movie S3 and fig. S2).

Because alcohol-treated males were more aggressive, we sought to determine if they would dominate sober males. We painted flies with a white paint dot to demarcate either the sober animal or the alcohol treated one (Fig. 2A). We determined whether the control fly or the PEA fly retreated and lunged. We also recorded the duration of interactions before retreats. For each pair of flies, we determined a Retreat Index (defined as [Number of Sober Fly Retreats – Number of PEA Fly Retreats]/Total Number of Retreats) and found that sober flies were much more likely to retreat compared to PEA flies (Fig. 2B). We also determined a Lunge Index (defined as [Number of PEA Fly Lunges – Number of Sober Fly Lunges]/Total Number of Lunges) for each pair and saw a slight but not significant probability that PEA flies were more likely to lunge (Fig. 2C). PEA flies also stayed on the food patch for longer than sober flies before retreating (Fig. 2D). We performed matched trials so half of the trials had the PEA flies marked with paint and the other half had the control flies marked with paint. In all, PEA flies were more likely to win a fight against sober flies (fig. S4).

**Figure 2.**
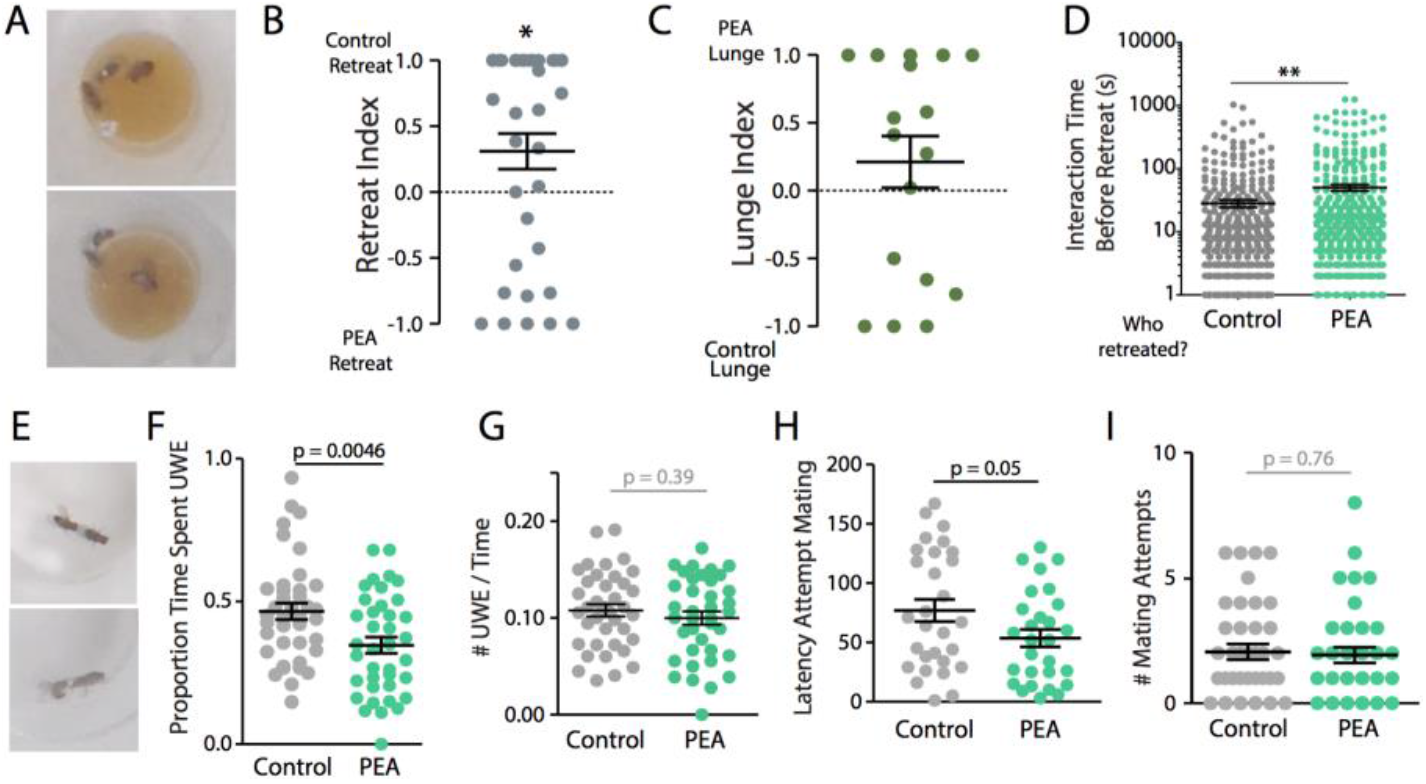
PEA flies are more dominant but are courtship defective. (**A**) Flies in aggression chambers with a white dot painted on their thorax. (**B**) Positive retreat Index values indicate sober flies are more likely to retreat, and negative values indicate PEA flies are more likely to retreat (one-sample *t*-test, μ_o_=0, *p<0.05, n=36). (**C**) Positive lunge Index values indicate that PEA flies are more likely to lunge, and vice versa for negative values (one-sample *t*-test, μ_o_=0, n=18 not significant). (**D**) Interaction time before retreat of a sober fly (gray) or PEA fly (green) (unpaired two-tailed *t*-test, **p<0.01, n=515-518) (**E**) Picture of courtship arena and a fly performing a unilateral wing extension (UWE) (top) and attempting to mate (bottom). (**F**) Proportion of time spent performing UWE, (**G**) number of UWEs, (**H**) latency to attempt mating (**I**) and number of mating attempts for each male fly tested (unpaired two-tailed *t*-test, n=38-39).

Sexual and aggressive behaviors are often intertwined on a neural and behavioral level and represent a major societal issue (*6*). We examined whether there were defects in courtship ability by video recording male flies in a small courtship arena in the presence of a virgin female. We quantified Unilateral Wing Extensions (UWEs) and mating attempts of the male fly (pictured Fig. 2E top and bottom, respectively). UWEs are important for successful copulation and occur when the male fly extends his wing to sing his courtship song to the female. PEA flies spent less time performing UWEs compared to control animals but performed roughly the same number of UWEs (Fig. 2F and G). In addition, PEA flies had a shorter average song bout duration than control animals (fig. S5). PEA flies were also faster to attempt copulation, but their total mating attempts did not change (Fig 2. H and I).

### *Fruitless* is a regulator of alcohol behaviors and is regulated by alcohol

The transcription factor gene *fruitless* (*fru*) has been implicated in sex-specific behaviors such as courtship, aggression, and alcohol preference (*7*–*9*). Fru^M^ has also been shown to regulate arborization patterns of neuronal circuits in a sex-specific manner (*10*). Previous work in our lab has demonstrated that FruM is necessary for sexually dimorphic preference for alcohol and for male flies to be able to acquire tolerance to alcohol (*9*). We hypothesized that alcohol-induced aggression could be regulated by this gene. The *fruitless* gene produces three alternative splice variants: the male variant, called *fru*^M^; the female variant, called *fru*^F^; and the common variant, called *fru*^COM^ that is expressed in both sexes (Fig. 3B). Transformer F (TraF) is a splicing regulator that suppresses alternative splicing events that generate the male-specific *fru*^M^ splice isoform (*11*) (Fig. 3C). Transgenic TraF expression suppresses production of *fru*^M^ and suppresses alcohol-induced aggression (Fig. 3A). However, TraF also alters splicing of transcripts from the *doublesex* (*dsx*) gene, which leads to expression of a female-specific *dsx* isoform known as *dsxF*. Overexpression of *dsxF* did not alter alcohol-induced aggression, indicating that *dsxF* does not modulate alcohol-induced aggression.

**Figure 3.**
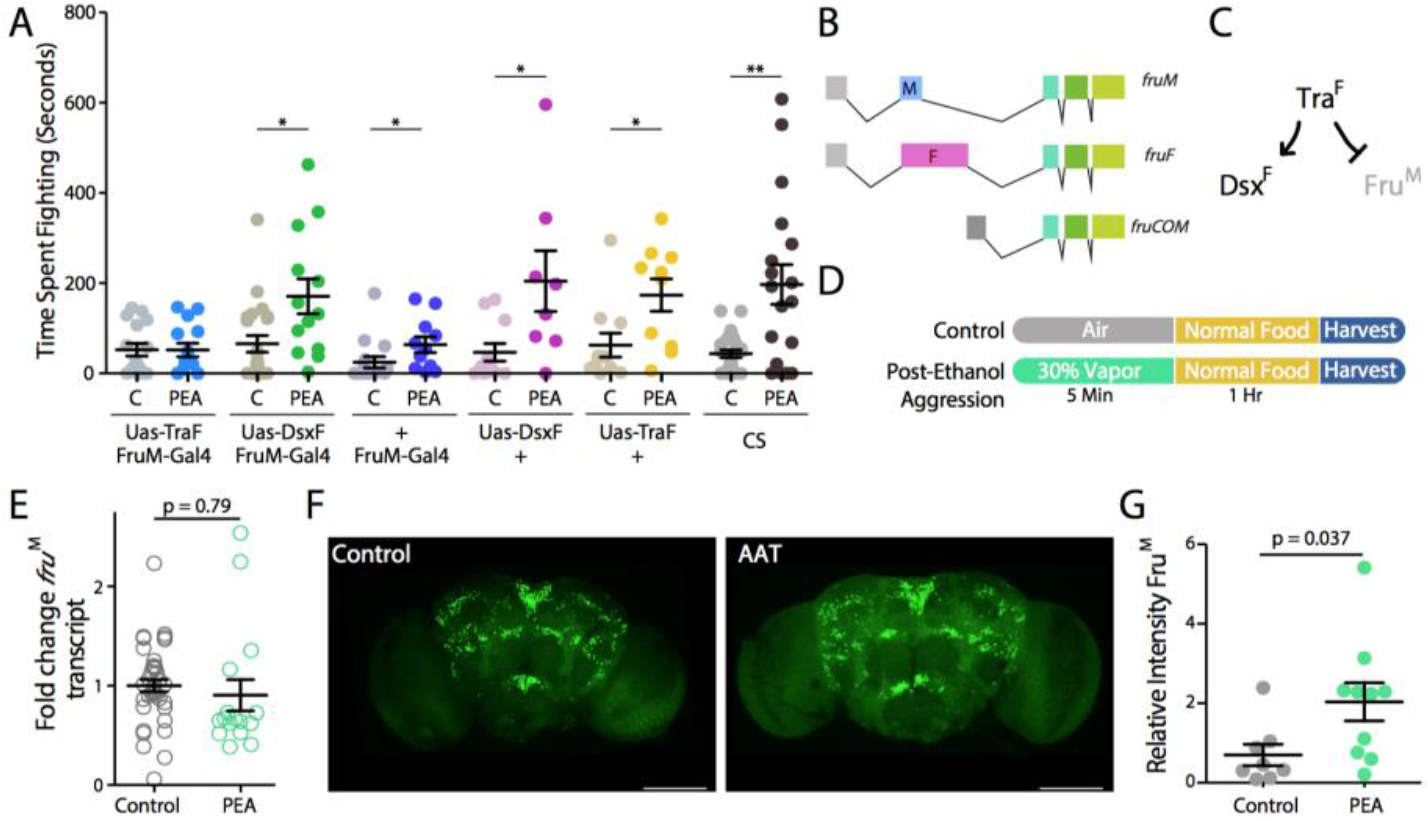
Fruitless is alcohol responsive and required for alcohol induced aggression. (**A**) Time spent fighting plotted across genotypes (± indicates no Uas or Gal4 transgene) and treatment (C = control, PEA = Post-Ethanol Aggression) (unpaired 2-tailed *t*-test, *p<0.05, **p<0.01, n=8-22). (**B**) Different fruitless splice isoforms, black arrows indicate primers used to target each splice isoform. (**C**) Transformer suppresses Fru^M^production and induces Dsx ^F^ expression. (**D**) Alcohol treatment paradigms for Post-Ethanol Aggression and Controls. (**E**) Fold change *fru*^M^ calculated using ΔΔCT method (two-tailed unpaired *t*-test). (**F**) Merged optical stacks across whole-brain volume for Control and PEA flies stained for Fru^M^. **(G)** Whole-brain relative intensity of Fru^M^stained fly brains. Relative intensity (normalized to control) of FruM signal (unpaired two-tailed *t*-test, n=8,10).

To determine if alcohol affects expression of *fru*^M^, we measured transcript-level changes in fly heads. PEA produced no significant changes in *fru*^M^ expression compared to air-treated controls (Fig. 3, D and E). Next, we measured Fru^M^ protein levels in the whole brain using a Fru^M^ antibody generously donated by Daisuke Yamamoto (*12*). PEA flies had increased Fru^M^ throughout the brain (Fig. 3F and G).

### Olfactory alcohol drives aggression

Within the Fru^M^-expressing circuit, Or67d-expressing Olfactory Sensory Neurons (Or67d OSNs) in the fly antennae are largely responsible for sensing the pheromone cis-vaccenyl acetate (cVa). When male flies smell cVa, they become more aggressive (*5*). These neurons contain the molecular architecture pictured in Fig. 5a. When a cVa molecule binds to LUSH, the cVa-bound LUSH displaces SNMP and activates Orco, depolarizing the neuron (*13*). There is also some evidence that cVa itself can bind to Or67d to activate the neuron independently of SNMP binding (*14*).

Previous work found that LUSH was involved in flies smelling alcohol. LUSH null mutants are unnaturally attracted to high concentrations of alcohol (*15*). More recently, LUSH was crystalized bound to ethanol, and the pocket shows conservation with ethanol-binding pockets of mammalian ion channels (*16*). Despite the link between alcohol and LUSH, no study has demonstrated a functional relationship between alcohol and cVa signaling. We hypothesized that olfactory exposure could also increase aggression in male flies by potentiating the cVa response.

To test this, we added alcohol into the fly food of the behavioral arena and then tested flies for aggression (pictured in Fig. 4A). To be sure that the alcohol used as an odorant did not also increase the blood alcohol level, we measured the BAC at the end of the 30-minute test period. This alcohol odor exposure paradigm did not produce a detectable increase in BAC (Fig. 4B). Flies that received an acute olfactory alcohol exposure (5% alcohol in the food) became significantly more aggressive (Fig. 4, C and D). Thus, acute olfactory exposure to alcohol increases aggression in the absence of an increase in systemic alcohol.

**Figure 4.**
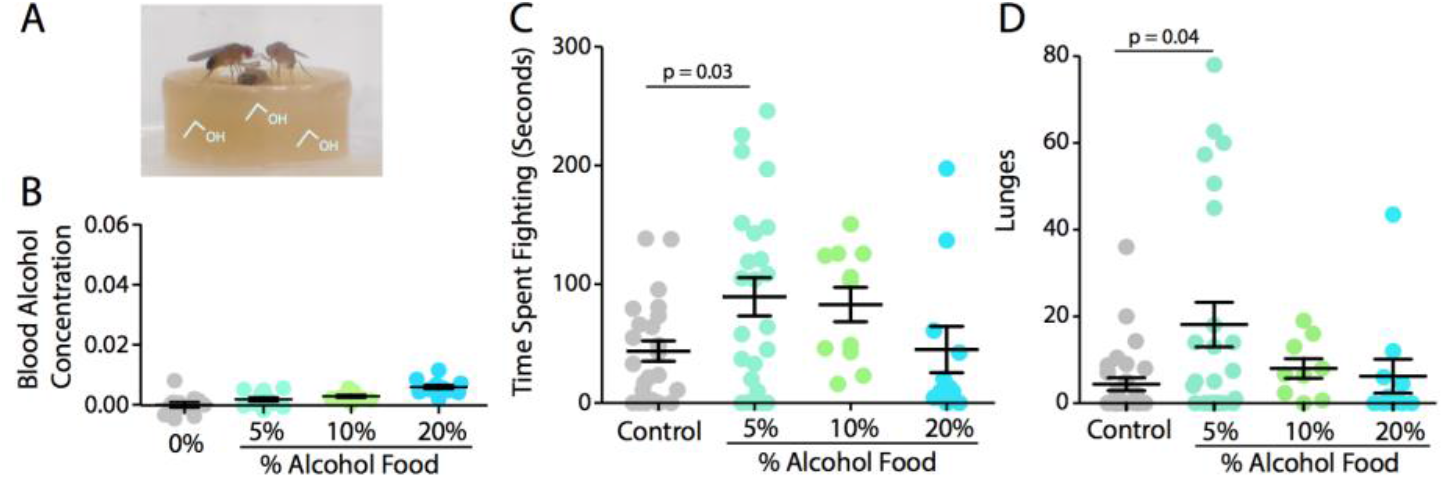
Alcohol odor potentiates aggression. (**A**) Picture of flies in aggression chamber with alcohol in fly food. (**B**) Blood alcohol concentration of flies exposed to alcohol food in aggression chamber. (**C**) Time spent fighting and (**D**) lunges for animals in aggression chamber with alcohol laden food (n = 11-24, One-way ANOVA with post-hoc Dunnett’s test).

**Figure 5.**
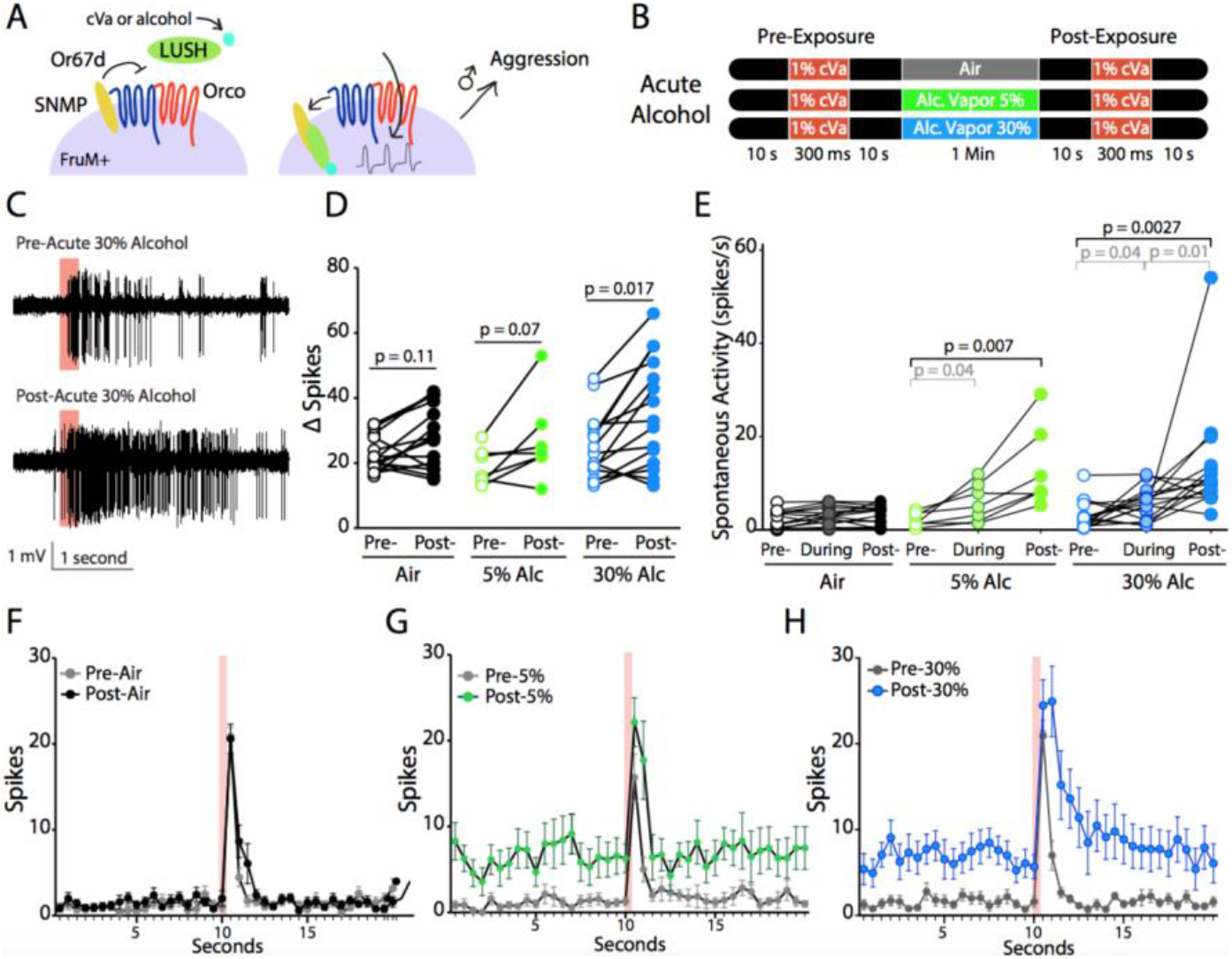
Acute alcohol potentiates the cVa response. (**A**) Overview of alcohol acting by two different mechanisms to promote aggression. (**B**) Acute alcohol treatment paradigms. (**C**) Raw spike traces taken using Autospike, red bars indicate when 1% cVa was applied. (**D**) ΔSpikes and (**E**) spontaneous activity plotted across treatment groups with lines drawn between different phases for each animal (unpaired 2-tailed *t*-test, *p<0.05, **p<0.01). Spikes were binned every 500 ms pre- and post-1% cVa application (red bar) for every animal (**F**) for air treated controls (n=15), (**G**) 30% acute alcohol treated animals (n=15), and (**H**) 5% acute alcohol treated animals (n=7).

Next, to determine if olfactory alcohol exposure electrophysiologically potentiates the cVa response, we performed in vivo single sensillum electrophysiology (SSR) of the T1 sensilla that contain the Or67d OSNs. For this paradigm, we delivered a 1% cVa puff, then administered air or alcohol vapor (30% or 5%), and then administered a second 1% cVa exposure (Fig. 5B). This concentration of 1% cVa was previously shown to be physiologically relevant as it mimicked the electrophysiological response of a male fly 1 cm away from a virgin female fly. From the recordings, we quantified spontaneous activity and evoked activity (ΔSpikes normalized to spontaneous activity; described in Methods). When we delivered acute 30% alcohol vapor, we observed a significant potentiation of the cVa response (Fig. 5, C and D). We found that an acute exposure to 30% and 5% alcohol increased spontaneous activity but that only an acute exposure to 30% alcohol increased evoked activity (Fig. 5, D and E and fig. S6).

Categorizing animals that either increased or decreased ΔSpikes, we find that the number of animals that had increased firing after alcohol was significant in both 30% alcohol-treated animals (13 increased, 2 decreased, χ^2^ test, p= 0.0045) and 5% alcohol-treated animals (5 increased, 2 decreased χ^2^ test, p= 0.0273) compared to air-treated controls (10 increased, 5 decreased χ^2^ test, p= 0.196). In addition, we looked for deactivation defects and found that flies treated with 30% acute alcohol had significant deactivation defects, whereas air-treated controls did not (Fig. 5, F-H and fig. S7). Taken together, these data demonstrate that acute exposure to alcohol potentiates the cVa response by increased evoked and spontaneous activity of the Or67d OSNs.

### PEA does not potentiate cVa

To assess whether PEA flies were also sensitized to cVa, we performed in vivo SSRs on flies treated with PEA and measured their response to 1% cVa (Fig. 6B and fig. S8). Compared to air-treated controls we saw no change in spontaneous activity, evoked activity, or deactivation changes (Fig. 6, C-E and fig. S9). Overall, PEA did not change the cVa response. Thus, the increase in aggression caused by PEA arises independently of the olfactory system and reflects a systemic effect of ethanol (Fig. 6A).

**Figure 6.**
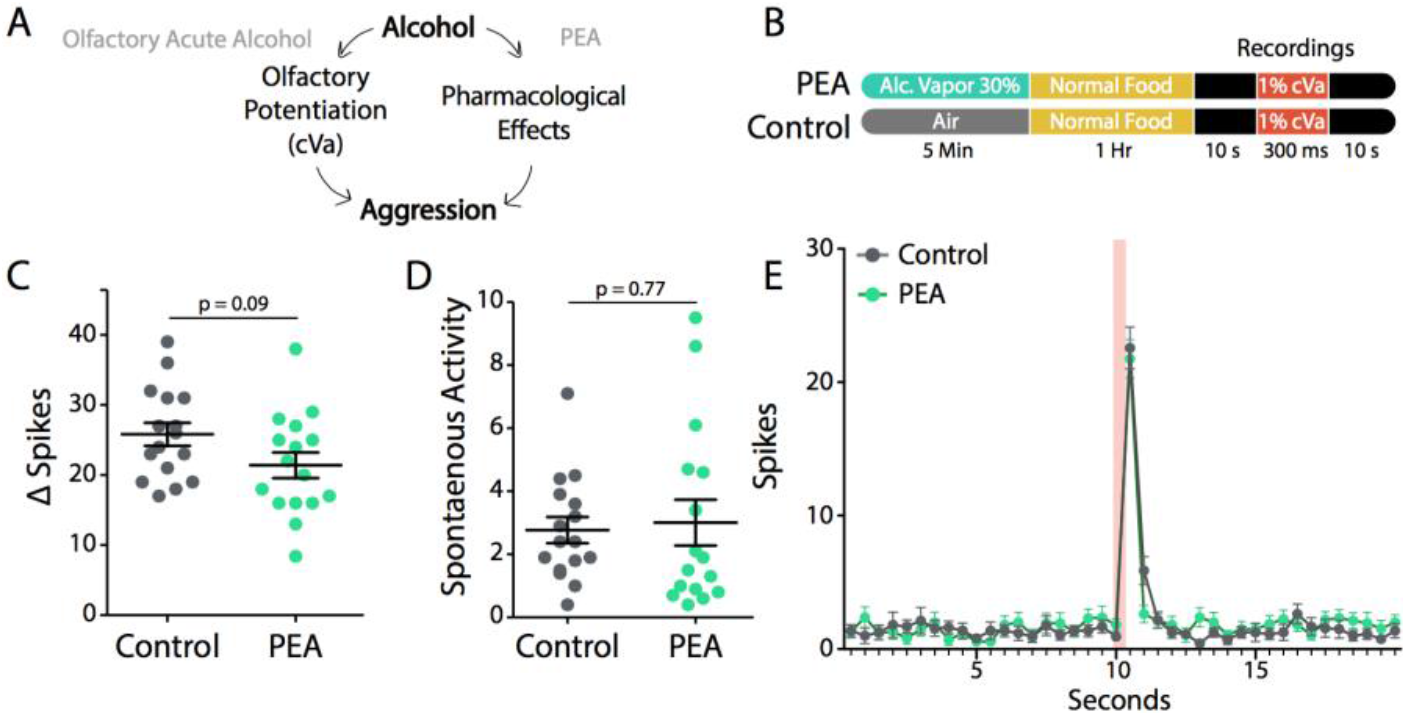
The cVa response is not potentiated after PEA. (**A**) The cVa transduction pathway in Or67d OSNs. (**B**) The PEA electrophysiology paradigm. (**C**) ΔSpikes and (**D**) spontaneous activity plotted for control (n=16) and PEA (n=16) animals (unpaired two-tailed *t*-test). (**E**) Spikes binned every 500 ms pre- and post-1% cVa treatment (red bar) for PEA vs. controls.

## Discussion

In this paper, we show that alcohol-induced aggression is an evolutionarily conserved behavior. We saw that low-dose alcohol induces aggression by two independent mechanisms—one that arises in response to the combined perception of the odor of alcohol and a male pheromone and one that originates from a systemic effect of alcohol in the CNS. Both avenues of action are likely relevant to the behavioral ecology of Drosophila.

We observed that low doses of alcohol act systemically to stimulate the expression of a transcription factor that defines male characteristics of the fly. The transcription factor is Fru^M^, which is the primary master regulatory switch that generates many male-specific behaviors. Expression of Fru^M^ is required for alcohol-induced aggression in male flies.

The mammalian orthologues of *fru*^M^ (DRSC/TRiP) are ZBTB45, 1, 39, and 24. The ZBTB (Zn finger and BTB protein-binding domain) family of transcription factors are known to regulate differentiation of immune cells, glia, neurons, and oligodendrocytes (*20*, *21*). ZBTB controls sex-related physiology such as spermatogonial stem cell renewal (ZBTB16), and the *SRY* gene contains a putative ZBTB binding site suggesting additional sex-specific regulation (*22*). The presence of a ZBTB transcription factor binding site in the SRY gene may mean that ZBTB activity directly modulates SRY expression. ZBTB also changes expression in response to alcohol in the mammalian brain (*23*, *24*). It is possible that ZBTB has a similar role in regulating alcohol-related behaviors in a sex-specific manner in mammals.

Furthermore, olfactory stimulation of aggression was also shown to occur because alcohol potentiates the signaling of cVA, a pheromone known to be responsible for male aggression. This mode of modulating aggression by alcohol is ethologically relevant. Drosophila in the wild are attracted to fermenting fruit as a preferred site for reproduction. The concentration of ethanol at these sites can be as high as 6% (*17*). In laboratory experiments, we have shown that females are attracted to alcohol (≤10%) food but that males are not (*9*). In addition, female flies preferentially oviposit in alcohol laden food because ethanol itself adds caloric value to the food substrate and can protect larvae from predators (*18*, *19*). In the wild, males may localize on alcohol-rich food because of the presence of females that are already attracted to these sites. Alcohol odors from these sites would then cause males to become more aggressive in an attempt to secure the female and the site for his own reproductive action. Thus, males would treat a female localized on a fermenting substrate as a higher value consort.

Currently, there is a lack of understanding of the neural and genetic mechanisms that regulate alcohol-induced aggression for a variety of reasons. First, aggression is a complex social behavior with a rich repertoire of actions, which makes it difficult to automate analysis. Automated analysis can not only expedite behavioral analysis but can also serve as an unbiased method for behavioral quantification that can reliably quantify behaviors not as obvious to the human eye. Second, performing large-scale screens is extremely difficult in organisms that do not have an accessible and extensive genetic toolkit. Lastly, most models of alcohol-induced aggression (especially in humans) test for accessory behaviors of physical aggression such as verbal aggression, non-physical punishment of a fictitious person (Taylor Aggression paradigm), and impulsivity; however, these types of paradigms crucially fail to accurately model alcohol-induced physical aggression (*25*, *26*). Identifying and characterizing alcohol-induced aggression in a model organism that rectifies these issues will inevitably allow us to better understand regulators of alcohol-induced aggression. Drosophila could serve to address these issues, as it has an extensive genetic toolkit, and behavioral paradigms that accurately model physical aggression, and some groups have already developed high-throughput automated analysis of aggression (*27*).

Taken all together, there are remarkable behavioral similarities between humans and flies when they are exposed to alcohol. Both show suppression of executive function by alcohol-induced aggression, and alcohol-induced promiscuity. Uncovering the neural and genetic substrates responsible for producing behavioral changes is essential for understanding the pathology of alcoholism.

## Acknowledgements

We thank A. Ghezzi, R. Maiya, E. Grantham, and M. Pomrenze for helpful discussions and technical input on the work; D. Yamamoto and R. Tanaka for the Fru^M^ antibody and technical assistance; A. Harris, R. Maiya, and J. Kirschman for feedback on the manuscript; the Bloomington Drosophila Stock Center for providing fly stocks.

## Funding

This work was supported by NIH Grants 2R01AA01803706A1 to N.S.A. and F31AA027160 and T32AA07471 to A.P. and NIH R01DC015230 to DPS and NIH 5T32GM008203 to EAS

## Author Contributions

A.P. designed the research and N.S.A supervised the work. A.P., T.T., L.G., and C.S. conducted and analyzed aggression behavior. A.P., J.P., and G.T. conducted and analyzed courtship experiments. A.P. and E.S. designed, conducted, and analyzed electrophysiology experiments and D.S. supervised the electrophysiology experiments. A.P. conducted the molecular biology and immunohistochemical staining. A.P. and N.S.A. interpreted the results and wrote the paper.

## Competing Interests

Authors declare no competing interests.

## Data and materials availability

All data are available in the manuscript or the supplementary materials; raw data and videos are available upon request.

## Materials and Methods

### Fly Handling

All flies were raised on standard cornmeal, molasses, and agar media in a 12:12 light:dark cycle. Flies used in aggression and courtship receptivity behavioral assays were all taken from group housed bottles as pupae and individually raised in vials. Flies used in imaging, immunohistochemistry, and qPCR were group housed.

### Alcohol Pretreatments

For all vapor treatments flies were pretreated with either 100% ethanol or 30% ethanol by volume in ddH_2_O. Vapor was administered at 2.5 L/min for 15 minutes for 100% ethanol and 5 minutes for 30% ethanol. Flies were transferred back into their food vials and allowed to recover.

We made 20% alcohol food by melting fly food in the microwave and waited until it cooled down to ~35° C and added 100% alcohol to reach 20% w/v alcohol. Flies were placed into fresh 20% alcohol food vials each day.

### Behavioral Tests

Aggression chambers were assembled using a fly vial cut 1 inch high and glued to one petri dish. The top of the chamber has 2 holes; 1 large hole is used for loading flies and one other smaller hole is in the center of the top and is used for circulation. Food wells were made by cutting 1.5 mL microfuge tube tops. Fly food was melted and pipetted into the microfuge tube tops. We added sucrose to the top of each fly food surface and a decapitated virgin female fly. Flies were loaded into the chamber by gentle aspiration and the video camera began recording 5 minutes after the flies were in the chamber. Aggression tests were conducted between the hours of 9 AM - 4 PM. Flies tested for aggression were between 4-6 days old.

Dominance was tested using the same aggression chambers. The video cameras were placed above the chambers to get an aerial view so that it was easier to determine the identity of the flies. Flies were painted with a small dot of white acrylic paint (Sherwin Williams Interior acrylic latex, 650428204) the day of eclosion and tested when they were between 4-6 days old. We video recorded for 30 minutes and videos were manually analyzed for when flies were both on the food patch, retreats from the fly food patch, and lunges as well as the identity of the fly.

Courtship behavior was tested in a plexiglass constructed chamber. We laser cut acrylic plexiglas into small circular arenas 1.5 cm across and 0.5 cm deep in a 3×4 array. We loaded chambers by gentle aspiration and video recorded flies for 3 minutes and manually analyzed videos for unilateral wing extensions (time of occurrence and duration) and attempts to copulate (time of occurrence). We tested alcohol treated males and sober males concomitantly. We used males 4-6 days old and virgin females 3-4 days old.

### RNA isolation and qPCR

Flies were CO_2_ sedated and collected between 1-2 days old and aged until they were 5 days old. Groups of 100-150 flies per N were flash frozen in liquid nitrogen and vortexed to separate heads from bodies and taken through a sieve to isolate heads. Fly heads were squished in β Mercaptoethanol containing squish buffer and isolated using a phenol-chloroform extraction. Primers for qPCR were designed using the IDT primer design online tool (idtdna.com). Reverse Transcription was performed using Superscript III Reverse Transcriptase (Invitrogen, Carlsbad, CA). We used ThermoFisher Power SYBR Green Master Mix (Waltham, MA, Catalog No. 4367659). We used a ThermoFisher Viia 7 Real-Time PCR System (Waltham, MA) with a Tm = 60 °C and 40 cycles per run. We used the ΔΔC_T_ method with a housekeeping gene (*cyp1*) and normalizing relative to control values.

### Primers *fruitless*

*fru*^M^ - FWD: 5’ CACAAGCGGAACATCGAAAC 3’ and

REV: 5’ AGGAAAATCGTCTCGAAGT 3’

RT: 5’ TTGTTGTTATCTGTGAGA 3’

*cyp1*- FWD: 5′-TCTGCGTATGTGTGGCTCAT-3′ and

*REV:* 5′-TACAGAACTCGCGCATTCAC-3′).

### Imaging and Immunohistochemistry

Flies were CO_2_ sedated before dissection. Brains were dissected in 1X PBS and fixed with 4% paraformaldehyde at 4° C for 1 hour. For Fru^M^ antibody staining the brains were washed in 0.2% PBT (1X PBS and 0.2% TritonX) three times for 15 minutes each wash, then blocked with 10% NGS for at least 3 hours. Brains were then transferred into guinea pig anti-Fru^M^ primary antibody in PBTN (0.2% PBT in 10% NGS) for at least 10 hours. Brains were then washed in PBT three times for 15 minutes each. We used Life Technologies Alex Fluor 488 goat anti-guinea pig IgG (Eugene Oregon). Secondary antibody was diluted in PBTN at 1:200. Brains were incubated atleast 12 hours with the secondary antibody at room temperature. Brains were then washed in 2 steps with PBT for 15 minutes each and dehydrated in glycerol in 3 one hour steps (10%, 50%, and 80% glycerol in PBS). Brains were mounted anterior side up on frosted Poly-L-lysine slides with Vectashield with DAPI (Vectorlabs, Burlingame, CA).

Brains were imaged with a Zeiss LSM 780 (Jena, Germany). We sampled at an interval of 1.81 μm at 20X through the whole volume of the brain. Gain was manually set for each group of brains analyzed and recorded. LSM files were analzyed using Metamorph Software (Molecular Devices, San Jose, CA).

### Ethanol Assay Kit

We used a Megazyme Ethanol Assay Kit Cat # K-ETOH (Megazyme, Bray, CO) to measure BACs. Flies were homogenized in ddH_2_O and centrifuged at 10,000 G for 10 minutes. The supernatant was taken and used to measure alcohol concentrations.

### Transgenic flies

Transgenic flies were ordered from the Bloomington Fly Stock Center at Indiana University (NIH P40OD018537), FruM-Gal4 UAS-GFP flies were generated by Rudolph Bohm (Texas A&M, Kingsville) and Or67d mutants or VainsC flies were generated and donated by Dean Smith (UT Southwestern) {Jin et al., 2008, #54861}. Full genotype for the FruM-Gal4 UAS-GFP flies are w-; sco/cyo; sb-fruP{64, 20XUAS,6XGFP}/TM2, Ubx. For Fru^M^ aggression experiments transheterozygous animals were generated by crossing w[*]; TI{GAL4}fru[GAL4.P1.D]/TM3, Sb[1] and P{w+mC=UAS-tra.F}20J7.

### Single Sensillum Recording Electrophysiology

Single Sensillum extracellular electrophysiology was conducted according to De Brunye et al. (1999). We used w^1118^ flies 3-5 days old because CantonS had higher variability in spontaneous activity and higher overall spontaneous activity that made data analysis difficult. Flies were assayed under a constant stream of charcoal filtered air (36 ml/min, 22-25°C) to prevent any contamination from environmental odors. cVa was diluted in paraffin oil (1% dilution); 1 μl was applied to filter paper and inserted into a Pasteur pipette; air was passed over the filter and presented as the stimulus. Signals were amplified 1000x, fed into a computer via a 16-bit analog-to-digital converter, and analyzed offline with AUTOSPIKE software (USB-IDAC system; Syntech). The low cut-off filter setting was 200 Hz and the high cut-off setting was 3 kHz. Action potentials were recorded by inserting a glass electrode in the base of a sensillum. Data analysis was performed as reported by Xu et al. (2). Signals were recorded starting 10-sec before odorant stimulation. cVA-evoked action potentials were counted by subtracting the number of spikes 1 sec before cVA stimulation from the spike number 1 sec after cVA stimulation (Spikes/sec). The recordings were performed from separate sensilla with a maximum of two sensilla recorded from any single fly.

Alcohol was delivered by adding it into the conical flask that feeds into the IDAC. The alcohol was made from 200 proof ethanol and diluted in ddH_2_O to make either 30% or 5% alcohol by volume (ABV) and always covered with parafilm. For acute treatments, the flow rate was roughly 36 ml/min, 22-25 °C and for PEA treatments we administered 30% alcohol at roughly 2.5 L /min.

**Spontaneous Activity** = Total number of spikes 10 seconds prior to cVa delivery / 10 seconds and **ΔSpikes** = Evoked activity (1 second during and after cVa delivery) – Spontaneous Activity.

**Fig. S1.**
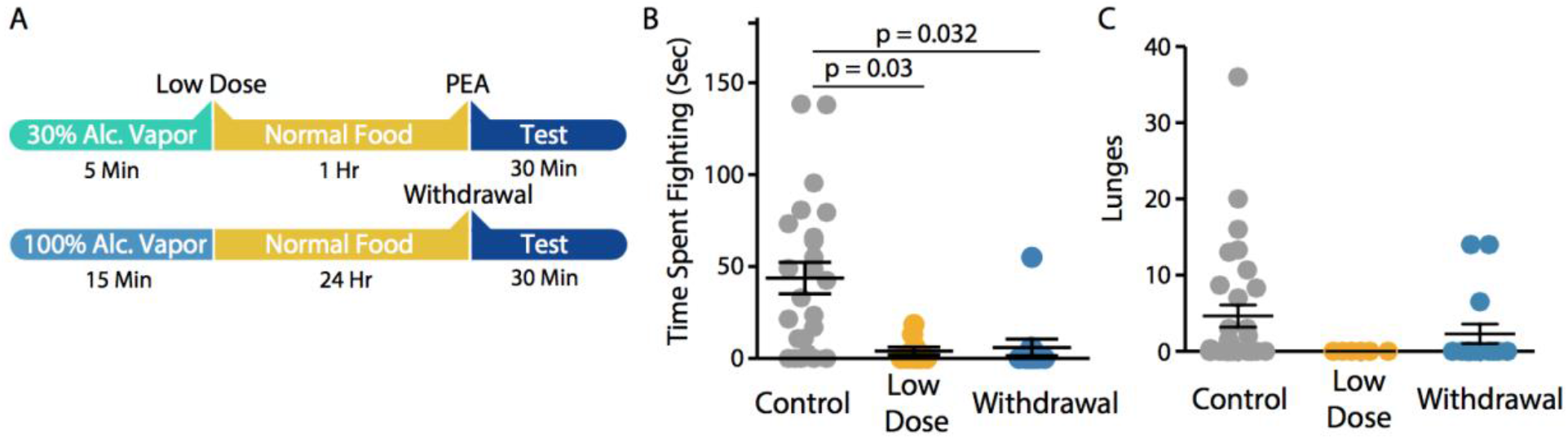
(**A**) Alcohol treatment paradigms for higher dose alcohol exposure (Low Dose and Withdrawal). (**B**) Both Low Dose and Withdrawal alcohol treatments suppressed time spent fighting and (**C**) lunges (unpaired 2-tailed *t*-test).

**Fig. S2.**
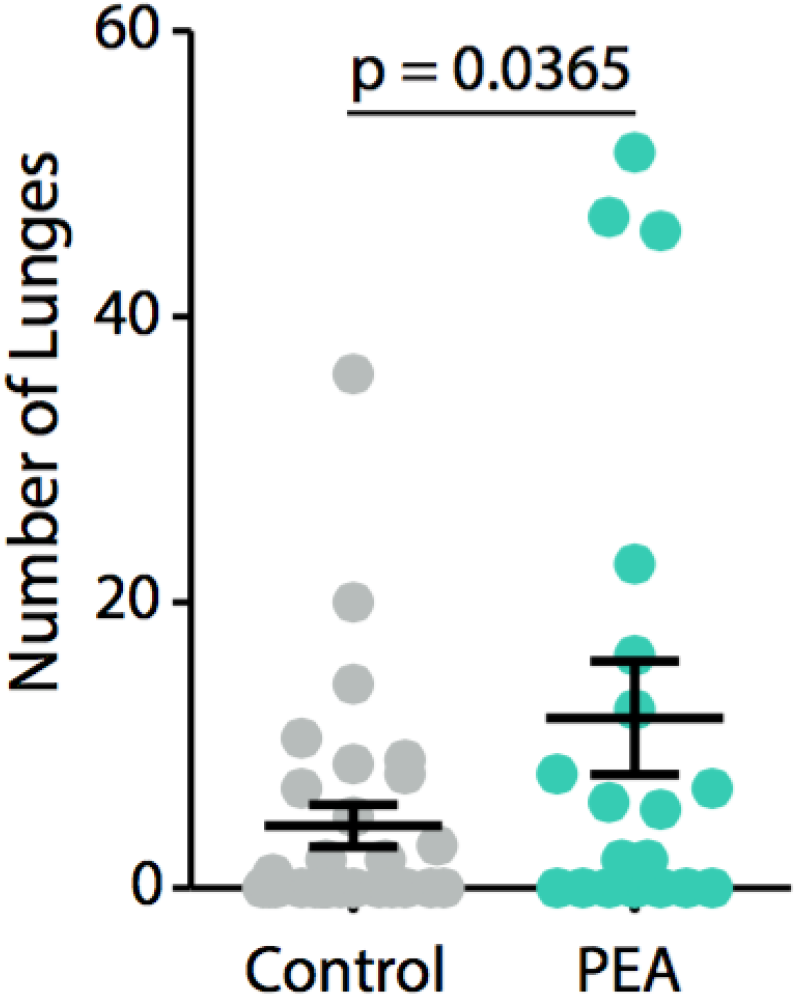
PEA flies lunge more than control animals. Unpaired two-tailed *t*-test.

**Fig. S3.**
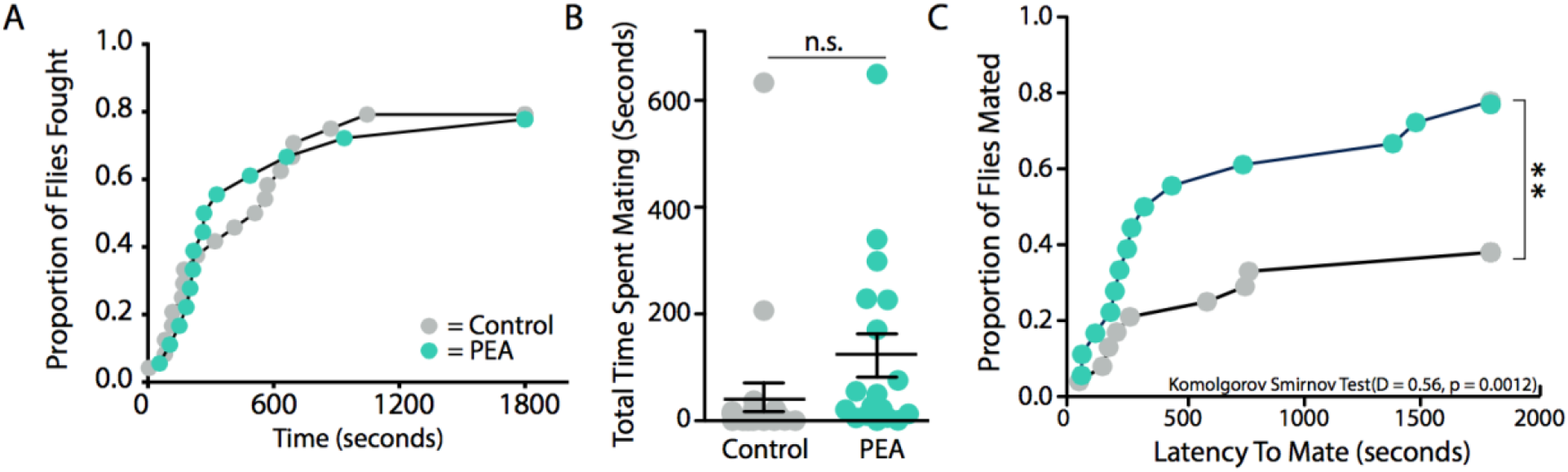
(**A**) There is no difference between PEA treated flies and Control animals in latency to fight or proportion of flies that engage in fighting. (**B**) There is no difference between the total time spent mating for either Control and PEA treated animals (n.s. = not significant, unpaired 2-tailed *t*-test). (**C**) The proportion of male flies that end up attempting to mate with the decapitated female by the end of the aggression assay is greater in the PEA treated animals (KS Test, p = 0.0012).

**Fig. S4.**
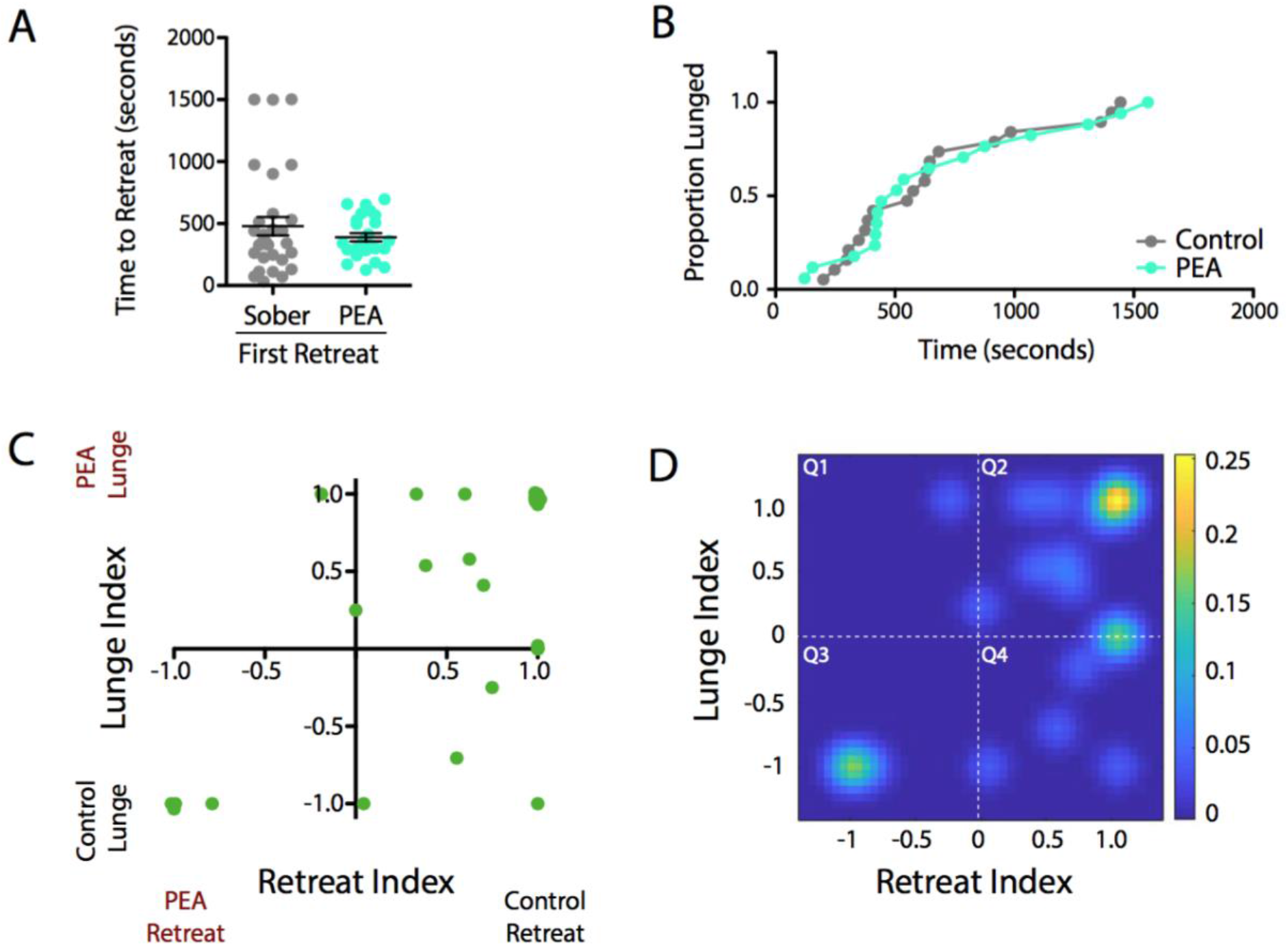
(**A**) There is no difference between retreat latencies of the first retreat between sober and PEA animals. (**B**) There is no difference between lunging latencies between sober and PEA animals. (**C**) and (**D**) Plots of Lunge Index vs. Retreat Index. More positive x-y values indicate greater chance of an PEA fly lunge and a sober fly retreat (PEA flies winning) and more negative x-y values indicate greater chance of sober flies lunging and PEA flies retreating (sober flies winning). PEA flies were less likely to retreat and clustered more in Q2 because they were also more likely to lunge. Sober flies that were more likely to lunge always became winners and less likely to retreat because almost no points were in Q1. However, even if PEA flies lunged less they would still be less likely to retreat (Q4). Heatmap generated in Matlab produced using binned data from (C).

**Fig. S5.**
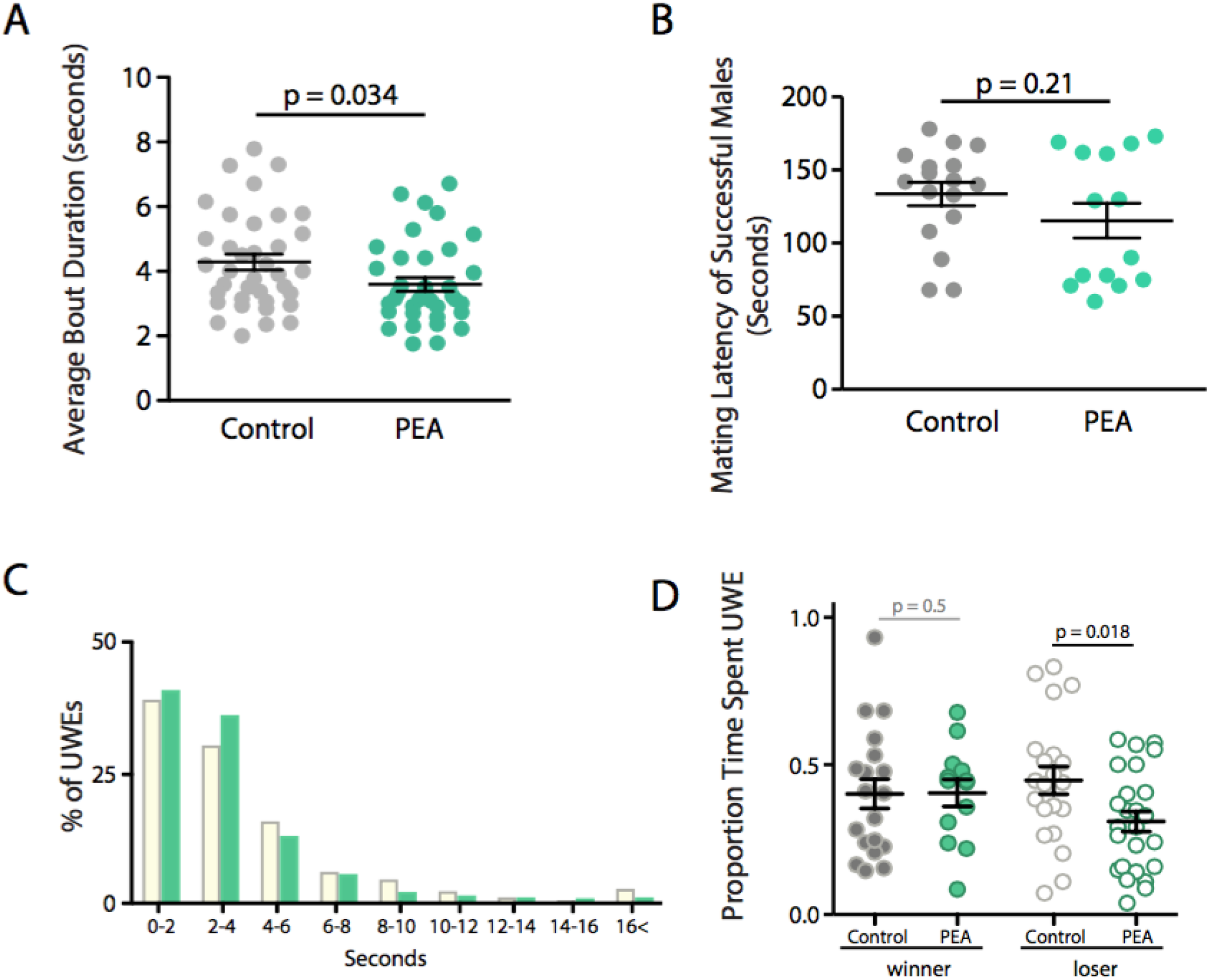
(**A**) PEA flies have a shorter average song bout duration than sober flies. (**B**) Mating Latency out of successful copulations does not differ significantly between sober and PEA animals. (**C**) UWE bouts binned in 2 second intervals and normalized to total number of UWEs per sober and PEA animals shows no change in the distribution of UWE bout lengths. (**D**) Grouped winners (mated males) and losers (unmated males) within sober and PEA animals shows that winners did not vary in their proportion of time spent performing UWEs, but losers did (unpaired two-tailed *t*-test).

**Fig. S6.**
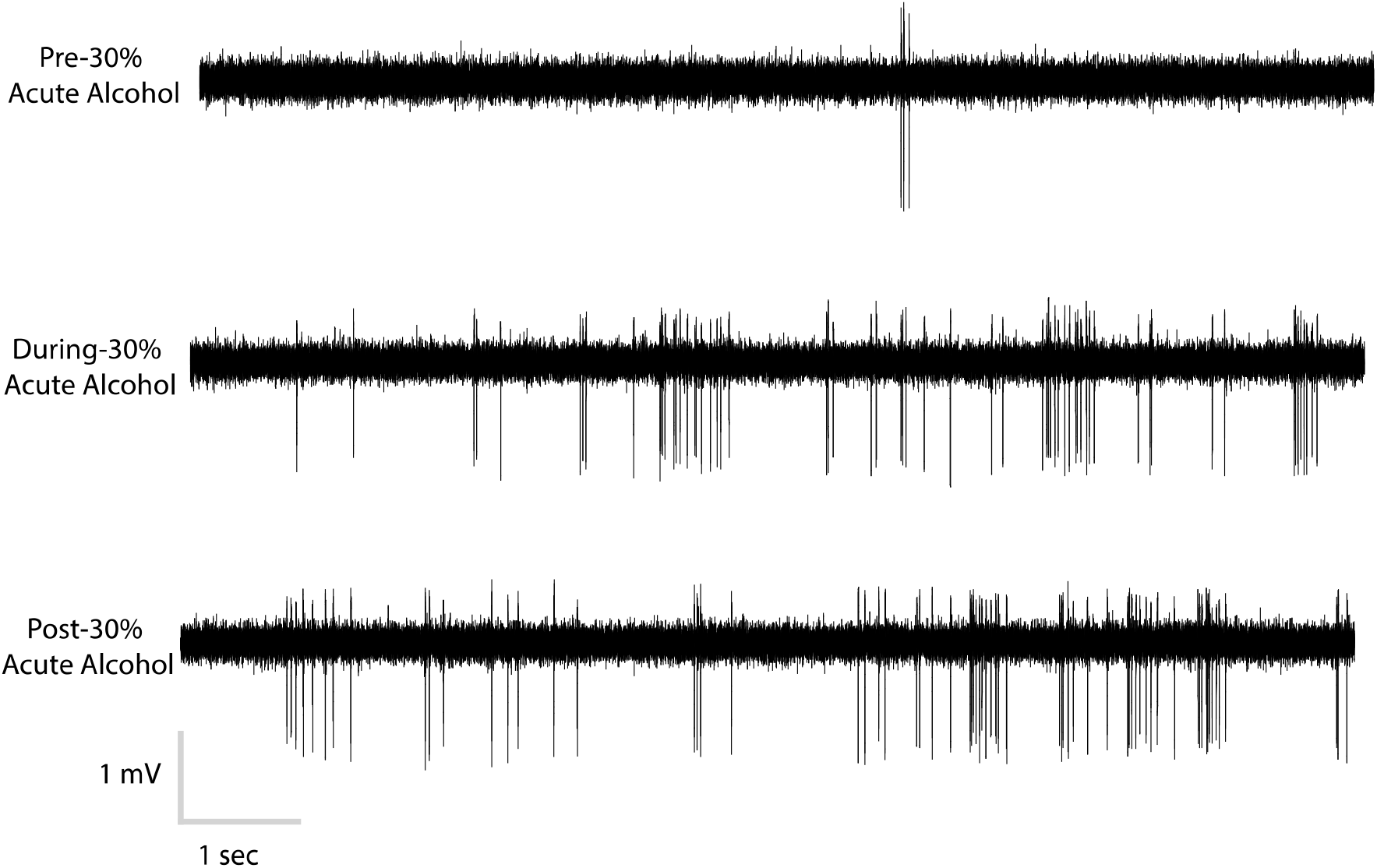
Sample traces of spontaneous activity before, during, and after 30% alcohol delivery pre-cVa delivery. Scale bars are 1 mV and 1 second.

**Fig. S7.**
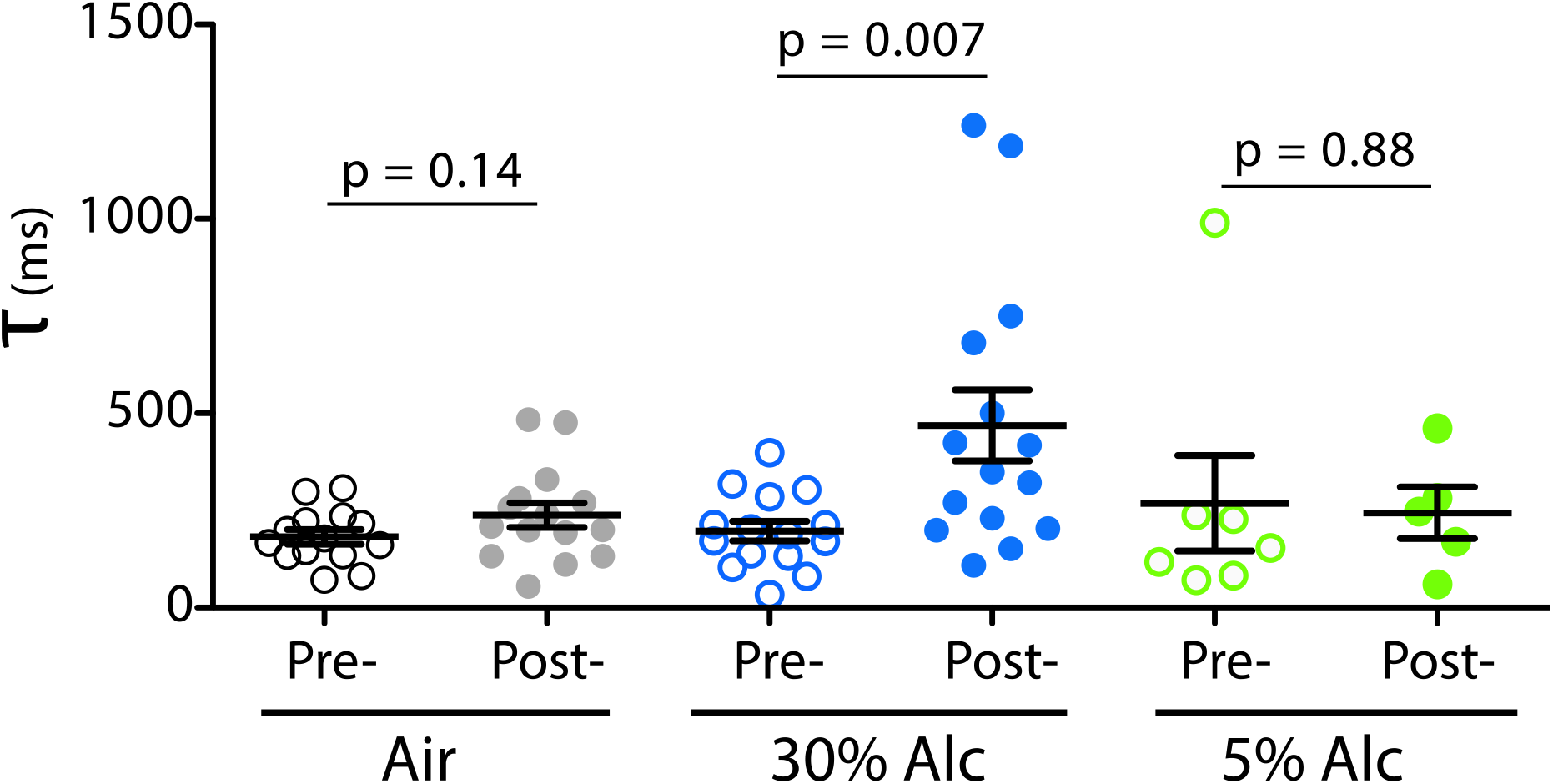
Time constants for each of the acute treatments pre- and post- alcohol or air. Time constant (τ) is calculated in ms from onset of peak activity and fit to a single exponential decay (unpaired 2-tailed t-test).

**Fig. S8.**
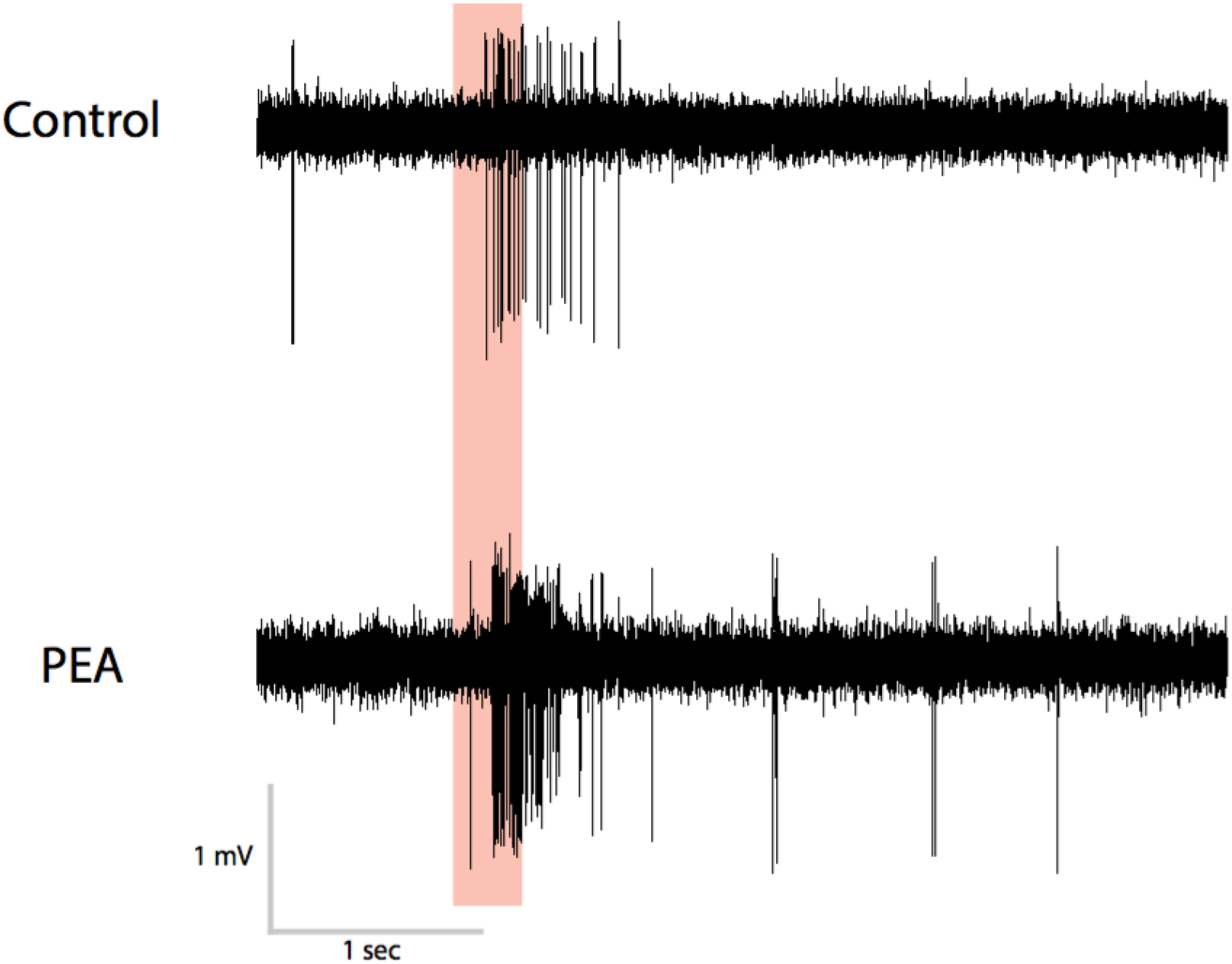
Sample traces of control and PEA animals. Red bar indicates when 1% cVa was applied. Scale bars are 1mV and 1 sec.

**Fig. S9.**
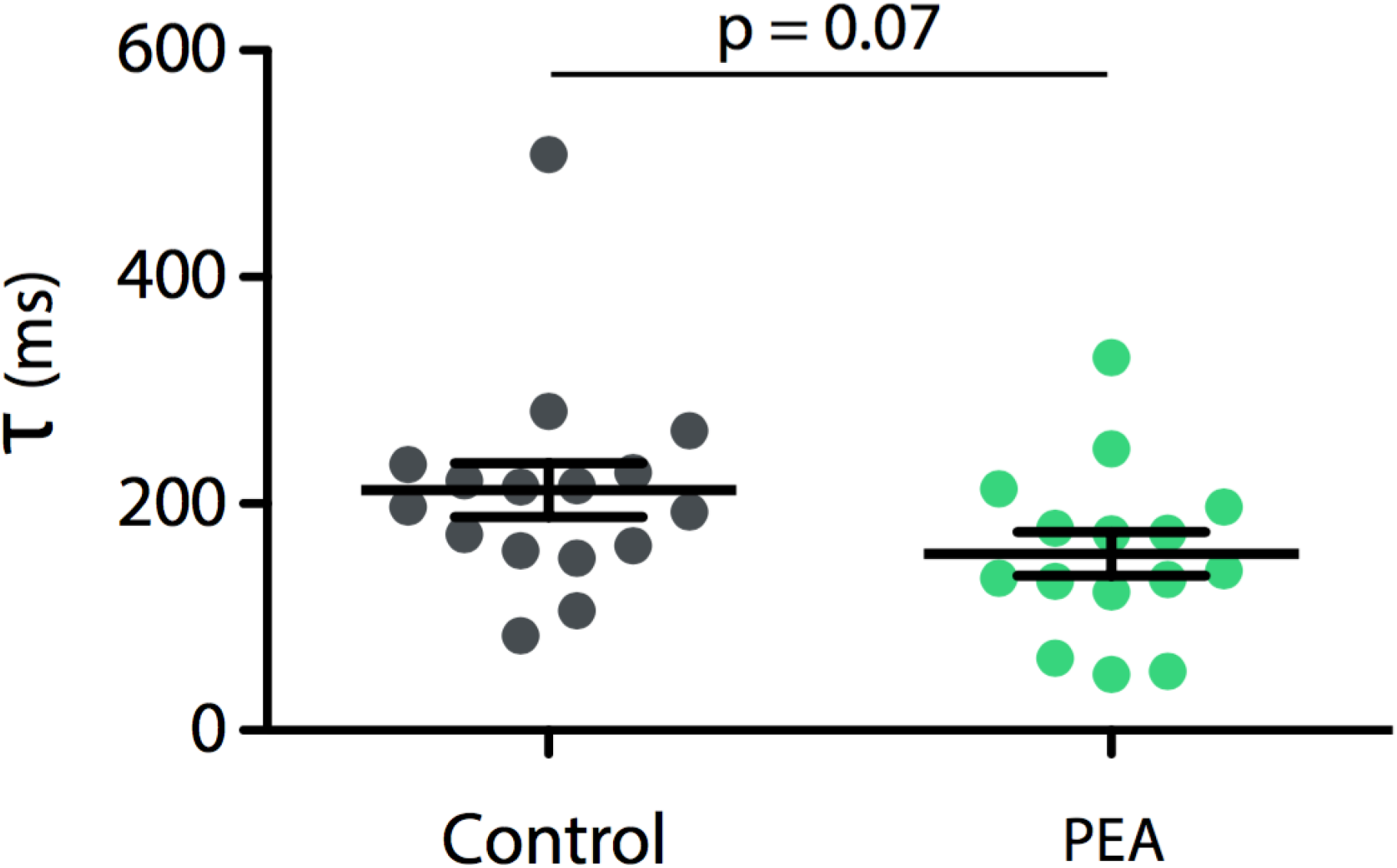
Time constants for each of the PEA treatment and control animals. Time constant (τ) is calculated in ms from onset of peak activity and fit to a single exponential decay (unpaired 2-tailed t-test).

**Movie S1.** Sober flies fighting.

**Movie S2.** PEA alcohol treated flies fighting.

**Movie S1.** Mating attempts with decapitated female.

